# Microplastics have shape- and polymer-dependent effects on soil processes

**DOI:** 10.1101/2020.06.02.130054

**Authors:** Anika Lehmann, Eva F. Leifheit, Maurice Gerdawischke, Matthias C. Rillig

**Affiliations:** Freie Universität Berlin, Institut für Biologie, 14195 Berlin, Germany; Berlin-Brandenburg Institute of Advanced Biodiversity Research, 14195 Berlin, Germany

**Keywords:** microplastic, soil aggregation, organic matter, shape

## Abstract

Microplastics are a diverse and ubiquitous contaminant, a global change driver with potential to alter ecosystem properties and processes. Microplastic-induced effects in soils are manifold as microplastics differ in a variety of properties among which the shape is of special interest. Microplastic shapes can resemble natural forms or be dissimilar from natural objects. Our knowledge is limited regarding the impact of various microplastic shapes on soil processes. Therefore, we conducted this two-part research comprising a meta-analysis on published literature and a lab experiment focusing on microplastic shapes- and polymer-induced effects on soil aggregation and organic matter decomposition. We here focus on fibers, films, foams and fragments as microplastic shapes.

In the meta-analysis, we revealed a strong research focus on fibrous and particulate microplastic materials, with films and foams neglected.

Our experiment showed that microplastic shapes are important modulators of responses in soil aggregation and organic matter decomposition. Fibers, irrespective of their chemistry, negatively affected the formation of aggregates. This supported the shape dissimilarity hypothesis. However, for other shapes like foams and fragments, the polymer identity is clearly an important factor co-modulating the soil responses.

Further research is needed to generate a data-driven foundation to build on our developing mechanistic understanding of the importance and consequences of microplastic shapes added to our soils.

## Introduction

Microplastic contamination of the environment is becoming an issue of increasing concern. Proposed as a new global change driver with increasingly realized impacts on marine, aquatic and terrestrial systems, microplastic-induced effects on ecosystem functions and processes are starting to be revealed, as research shifts from a more ecotoxicological focus to more fully embrace an ecosystem perspective (Rillig and Lehmann, 2020). But microplastics are not a monolithic issue and neither are the effects they induce. Microplastics are a group of synthetic polymers encompassing a wide diversity in sizes, shapes, chemistries and additives (Wagner et al., 2014). In addition, plastic particles are exposed to weathering in the environment, further enhancing this diversity in properties: microplastics are either produced as primary microplastics or secondarily via degradation in the environment to fragments < 5mm (Hartmann et al., 2019). Our knowledge about the effects of microplastics on soils is still limited, although they likely are exposed to microplastic broadly and at increasing levels, especially agricultural and urban soils. Microplastics enter soils via atmospheric deposition (Allen et al., 2019), plastic mulching (Steinmetz et al., 2016), sewage sludge (Nizzetto et al., 2016) and compost (Weithmann et al., 2018). Microplastics are ubiquitous and persistent contaminants. After entering the soil matrix (Rillig et al., 2017), they have the potential to alter soil properties and processes.

Microplastics can affect soils as a function of their chemistry, but also through their shape (Rillig et al., 2019). Microplastics can resemble natural shapes, or they can be quite dissimilar from naturally-occurring objects in soil. In soils, microplastic beads and fragments can be almost indistinguishable from e.g. sand grains. Microplastic fibers, rigid and unbranched filaments, diverge clearly from hyphae, plant roots or other natural filamentous, linear organisms or structures. Other microplastic shapes like films and foams have no analogs in a natural system due to their properties and size. Rillig at al. (2019) highlighted the importance of shapes in the context of the population of naturally occurring particles at any given scale, suggesting that shapes more dissimilar from naturally occurring particles might cause stronger effects than plastic particles that resemble natural shapes. Research on terrestrial systems revealed that a key feature of soil - its structure - is often negatively affected by microplastic fibers, while for microplastic beads and fragments there is a range of positive to negative effects (e.g. Machado et al., 2018; Boots et al., 2019; Machado et al., 2019; Zhang et al., 2019). The spatial arrangement of primary particles into aggregates, and the resulting pore networks, together strongly control soil processes, including biogeochemical cycles and soil carbon storage and processing. For example, soil organic carbon can be physically protected inside of soil aggregates. The protection of soil organic carbon can be affected by microplastic indirectly via effects on soil structure or via effects on microbial activity, but observed effects are variable, shifting with microplastic concentration and properties (Liu et al., 2017; Machado et al., 2018; Boots et al., 2019). To better understand effects of microplastics on soil aggregation and organic matter processing, it is thus necessary to more systematically examine key microplastic properties, for example their shape.

To this end, we here present a two-part study, consisting of a quantitative synthesis based on available data, and a laboratory experiment systematically varying microplastic shapes. Research on microplastic-induced effects on soil systems is accumulating and with it the need for a quantitative evaluation of the consequences for soil structure and decomposition. Hence, we here conducted a meta-analysis on published articles presenting microplastic-induced effects on soil aggregation and soil organic matter loss. We identified overall 9 articles for which we extracted data on soil aggregate formation, stability and soil organic matter loss and information on microplastic shape and polymer. With the resulting datasets we investigated (1) the overall impact of microplastic on soil aggregate formation and stability and soil organic matter loss, irrespective of shape and polymer, and (2) how the effects are modulated by shape and polymer.

The biggest limitation of available research on microplastic induced effects on soils is the focus on just filamentous and particulate shapes. To add new insights on the importance of different shapes we designed a laboratory study in which we tested the effect of four different shapes, each represented by three different polymers, on soil aggregation and organic matter decomposition. We hypothesize (1) that fibers, films and foams will affect soil structure and organic matter decomposition more strongly than fragments due to their shapes which are more dissimilar to natural shapes present in the soil; and (2) that the shape is more important than the polymer.

## Material and methods

### Meta-analysis

First, we conducted a meta-analysis on published articles reporting microplastic effects on soil aggregation and/ or soil carbon fractions. We followed the PRISMA guidelines (for further information see supplementary information; Fig. S1).

### Literature search

We conducted a three-step literature search on 2^nd^ of April 2020 to identify and collect articles for our database. First, we used the two search strings “(*microplastic* AND soil*)” and “(((*microplastic* OR microfiber*\*) *AND* (*decompos* OR* “*organic matter*” *OR litter*)) *NOT* (*marine OR sea OR water OR aquatic*))” to run topic searches in Web of Knowledge (Thomson Reuters) focusing on articles in English language. Both search outcomes were combined with the “OR” option to generate one search outcome with 276 hits eliminating duplicates (search 1: 174 hits, search 2: 120 hits). These articles were screened by title and abstract to identify potential candidate articles that were then further investigated according to our inclusion and exclusion criteria. (1) Articles had to present data on soil aggregation (e.g. water-stability, size class weights) and/ or decomposition (i.e. mass loss, rate of substrate degradation) of organic material (e.g. litter, cellulose stripes) for microplastic contaminated samples and associated controls. (2) Only soil based systems were applicable. (3) The microplastic products (< 5 mm; Hartmann et al., 2019) had to be incorporated into the test soil. Foils, mulches or other types of plastic layers on top of the soil were not taken into account. (4) Biodegradable plastics (e.g. biodegradable polylactic acid) were excluded since they could be degraded during the course of the experiment and since they are currently not widely used. (5) Experiments without artificial organic matter application or additional stressors (e.g. drought) were chosen. (6) In cases where data are presented in presence/absence of plants, we preferred no plant data, to avoid confounding effects of plant roots on soil. (7) We chose data for the highest presented microplastic level and last harvest day.

In the second search step, the reference and citation lists of all matches were screened for further matching articles. In the final step, we searched Google Scholar and the preprint server bioRxiv and EcoEvoRxiv to capture recent publications that were not listed by Web of Knowledge at the time of the search.

The complete search yielded 8 and 1 articles, respectively, for the soil aggregation and decomposition focused search runs. With these articles we build three datasets: the (1) soil aggregate formation, (2) stabilization and (3) decomposition dataset.

Soil aggregation data was presented for wet- or dry-sieving techniques. For both techniques, the mean weight diameter (MWD) was the dominant metric giving information for aggregate stability for multiple aggregate fractions of decreasing size. For wet-sieving, a test focusing on one soil fraction >4mm was also available (i.e. water-stable aggregates <4mm [WSA]). For the soil aggregate stabilization dataset, we preferred MWD over WSA due to higher resolution of aggregate size class contributions. We either used presented MWD metrics or calculated it following the formula: 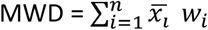, where 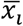 is the mean diameter of each aggregate size fraction and *w_i_* is the proportion of the aggregate mass in each size fraction and the overall sample weight.

For the aggregate formation dataset, we included only data for aggregate fractions larger than the soil used at the beginning of the experiment; i.e. in cases where the soil was sieved to 2 mm prior to the experiment, we only considered data on aggregate weights in soil fraction >2 mm at the end of the experiment. This fraction was compared to the overall sample weight to calculate the percentage of newly formed aggregates (NFA): NFA= size class _> *initial size*/_ sample weight _*overall*_.

For the organic matter loss dataset, we collected data only on decomposition of organic material added to the test system by the experimenter and whose degradation was subsequently assessed. Results of organic matter dynamics (e.g. whole soil or aggregate associated carbon) were not suitable: the applied microplastic itself is a source of carbon, which adds to the soil carbon pool (Rillig, 2018) and which is not distinguishable from other carbon-containing organic molecules with standard methods.

### Effect sizes

For the three datasets focusing on soil aggregate formation, stabilization and soil organic matter loss, we calculated specific effect sizes using the natural log response ratio of the treatments and control groups following the function: log response ratio = log(X_T_/X_C_). X_T_ represents the mean data of microplastic treated samples and X_C_ the corresponding microplastic-free control means. The effect sizes were calculated with the function escalc() in the package “metafor” (Viechtbauer, 2010) by integrating treatment and control means, the associated standard deviation (sd) and number of replicates of experimental units (*n*). In cases where only the standard error (se) was reported, we calculated sd as follows: sd= sqrt(n) x se. One article did not report any metric of variance, here, we used the median of all calculated effect size variances as a surrogate (Copas and Shi, 2000).

### Moderator variables

We collected data on the shape and polymer of the tested microplastics. The variable “shape” comprised fiber and particles; particles included all non-linear shapes e.g. fragments, beads and spheres. In the variable polymer we listed all tested microplastic polymers. In one case a mixture of two fibers (polyester and polyacryl) was tested; we included this study and labeled it as “mix” in the variable “polymer”.

### Statistics

All statistics were conducted in R version 3.6.1 (R Development Core Team, 2014). The constructed datasets comprised multiple effect size values per study. This was e.g. due to the testing of multiple microplastic types and polymers in one experiment. In only one case it was necessary to use an effect size merging approach because individual effect sizes (presented for different fungal strains) did not contribute to our moderator variables. In this case, we applied a phylogenetically corrected merging by using the rma.mv() function in the “metafor” package (Viechtbauer, 2010; Anderson, 2016). The model included a phylogenetic tree as a random factor (for tree construction please refer to (Lehmann et al., 2019)).

The final datasets were analyzed by a random-effects model with the function rma.uni() in the R package “metafor” implementing study weighting by the inverse of the effect size variance and the restricted maximum likelihood approach. Data were tested for normal distribution via qq-norm plots. As sensitivity analyses, we applied funnel plots and trimfill() functions in the R package “metafor”. The outcomes of these tests can be found in the supplementary information (Fig. S2).

### Experiment

We conducted a laboratory experiment in a soil system to test for effects of a diverse set of microplastic polymers representing different shapes. The experiment comprised four shapes, each with three polymers, resulting in 12 microplastic treatment levels and a plastic-free control. For each microplastic level we had 10 and for the controls 40 replicates, yielding overall 160 experimental units.

### Microplastic materials

The plastic treatment consisted of “simulated” secondary microplastic products (in the sense that they were not produced in the environment), which were processed for this experiment from commercially available products (Fig. 1; for details see Table S1). For microfibers, we chose polyamide (nylon), polyester and polypropylene obtained by manually cutting ropes. Microplastic films were produced by a hole puncher and manually cutting polyethylene terephthalate, casted polypropylene and polyethylene foils from supermarkets and gardening supplies. For microplastic foams, we used polyethylene, expandable polypropylene and polyurethane products from packaging and foam rubber padding materials. Microfragments were produced from polycarbonate (CDs/ DVDs), polyethylene terephthalate (water bottles) and polypropylene (planting pots). The microplastic foams and fragments were generated via blending (Philips Pro Blend 6 RD). The microfibers and –films could not be obtained by blending due to their high flexibility, not even after freezing the products with liquid nitrogen.

**Fig. 1.**
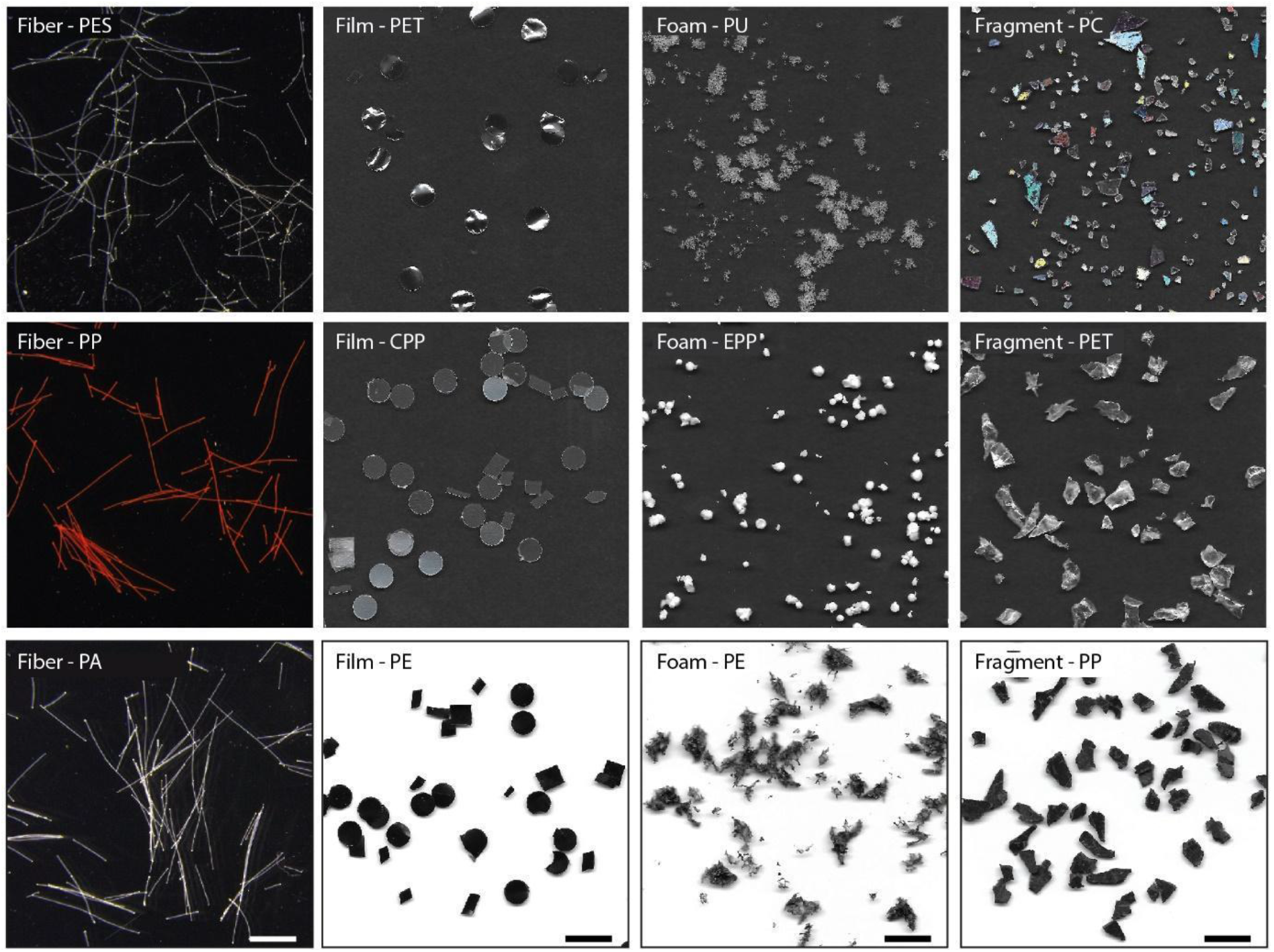
Overview of used plastic types (fibers, films, foams and fragments) and the respective polymers: PES = polyester, PP = polypropylene, PA = polyamide, PET = polyethylene terephthalate, CPP = casted polypropylene, PE = polyethylene, PU = polyurethane, EPP = expandable polypropylene, PC = polycarbonate. The white scale bar responds to the fibers only and represents 1mm while the black scale bars for films, foams and fragments represent 1cm.

The secondary microplastics varied in their size range but were all < 5 mm, and thus met the definition for microplastics (Table 1; Hartmann et al., 2019). The materials were surface sterilized (microwaving for 2 min, 630 watts) before introducing them into the test systems, all at a concentration of 0.4% (w:w).

**Table 1.** Length and width/ diameter distribution for the used plastic shapes (fibers, films, foams and fragments) and the respective polymers: PES = polyester, PP = polypropylene, PA = polyamide, PET = polyethylene terephthalate, CPP = casted polypropylene, PE = polyethylene, PU = polyurethane, EPP = expandable polypropylene, PC = polycarbonate.

| Shape | Polymer | Length |  |  | Width/diameter |  |
| --- | --- | --- | --- | --- | --- | --- |
|  |  | N | mean | sd | mean | sd |
| Fiber | PES | 100 | 0.899 | 0.736 | 0.030 | 0.004 |
|  | PP | 100 | 1.325 | 0.697 | 0.028 | 0.004 |
|  | PA | 100 | 1.248 | 0.794 | 0.030 | 0.006 |
| Film | CPP | 100 | 5.378 | 1.913 | 4.018 | 1.579 |
|  | PE | 100 | 4.478 | 1.087 | 3.208 | 1.623 |
|  | PET | 100 | 5.086 | 0.833 | 4.289 | 1.469 |
| Foam | PU | 100 | 2.858 | 2.203 | 1.691 | 0.975 |
|  | PE | 100 | 4.565 | 3.116 | 2.133 | 1.482 |
|  | EPP | 100 | - | - | 2.039 | 0.802 |
| Fragment | PP | 100 | 4.717 | 0.412 | 2.693 | 0.880 |
|  | PC | 100 | 2.065 | 1.183 | 1.157 | 0.642 |
|  | PET | 100 | 4.939 | 2.401 | 2.621 | 1.098 |

### Test system

We used 6cm Petri dishes filled with 15 g of an unsterilized local soil sieved to <2 mm particle size. With this test system, we focused on soil-derived responses related to structural changes and organic matter decomposition. The soil had a sandy loam texture (Albic Luvisol following FAO classification; 74% sand, 18% silt, 8% clay), with 6.9 mg/100 g P (calcium-acetate-lactate), 5.0 mg/100 g K (calcium-acetate-lactate), 0.12% total N and 1.87% total C content and a soil pH of 7.1 (Rillig et al., 2010). The soil for each system was mixed individually with the designated microplastic material, and the controls were mixed with the same energy and for the same time to apply an equivalent disturbance.

In each system, we inserted a miniature tea bag (Keuskamp et al., 2013) made of 2.5 x 5cm rectangles of 30 μm nylon mesh (Sefar Nitex) and an impulse sealer (Mercier Corporation, product no. 127174). The tea bags were filled with 300 mg green tea composed of 49 % C (Lipton green tea, Sencha Exclusive Selection, (Keuskamp et al., 2013)). The tea had a C:N ratio of 12, which can be considered as high quality litter that is easy to decompose. The test system was initiated by adding autoclaved (121°C for 20min) distilled water to reach a water holding capacity of 60%. Each unit was sealed with parafilm and stored in the dark at 22°C for 4 weeks. At harvest, the test systems were opened, dried at 40°C and stored until further usage.

### Measurements

We evaluated three system responses to the microplastic contamination. (1) new formed aggregates >2.0 mm, (2) the percentage of water-stable aggregates and (3) organic matter loss in the miniature tea bags. The soil samples were carefully removed from the Petri dishes and passed through a 4 mm sieve to break up the dried aggregates, and we retrieved the tea bag.

Subsequently, we tested the samples for the amount of aggregates formed *de novo* from the initial soil <2.0 mm after 4 weeks of incubation. For this, we placed the prepared soil fraction <4 mm on a second 2 mm mesh which we moved vertically two times to allow separation of the soil sample into the two fractions >2 mm and <2 mm while avoiding abrasion. The amount of newly formed aggregates was then standardized by the overall weight of the sample:

Newly formed aggregates >2 mm (%) = (fraction_> 2 mm_ / [fraction_>2 mm_ + fraction_<2 mm_]) × 100.

In the next step, we carefully mixed the separated fractions, placed 4.0 g of each sample on a small sieve with 250 μm mesh size, which were allowed to capillarily re-wet with distilled water before inserted into a sieving machine (Agrisearch Equipment, Eijkelkamp, Giesbeek, Netherlands). The machine sieved for 3 min in water-filled tins filled with distilled water (Kemper and Rosenau, 1986). By this technique, we separated each sample into an unstable fraction, and a water-stable fraction with a size > 250 μm. The water-stable fraction was further treated to extract sand particles and organic debris. The weight of starting soil volume (4.0 g) and the water-stable fraction were corrected for this coarse matter fraction.

Percentage of water-stable aggregates (%) = [(water stable fraction - coarse matter) / (4.0 g - coarse matter)] x 100.

Finally, we determined organic matter loss from the miniature tea bags by measuring the relative mass remaining (g g-1) on a dry matter basis. Nine tea bags were lost during the harvest.

### Statistics

All analyses were conducted in R (R Development Core Team, 2014). We tested the effect of our microplastic treatment (three polymers per four shapes) on the three soil-derived responses in a two-step approach. In the first step, we used an estimation method to generate unpaired mean differences (treatment minus control) via bootstrapping (5000 iterations) implemented in the R package “dabestr” (Ho et al., 2019). This approach focuses on the magnitude and the precision of an effect further supported by the sampling-error distribution which is also plotted. It allows clear visualization and identification of positive, negative and neutral effects of treatments compared to the controls. If the unpaired mean difference confidence interval (CI) overlaps the zero line (line of no effect) the corresponding effect is neutral, i.e. the treatment caused no detectable response compared to the control.

In the second step, we evaluated our findings by generalized least square models in the “nlme” package (Pinheiro et al., 2018). We accounted for heterogeneity in the variables shape and polymer by applying the varIdDent() function. We checked model residuals for heteroskedasticity and normal distribution. For multiple comparison tests of the microplastic treatments against the control samples, we used the glht() function in the package “multcomp” (Hothorn et al., 2008) to run a many-to-one comparison procedure (Dunnett, 1955). We tested the hypothesis that the mean difference of the microplastic treatments and the controls is ≥ zero (treatment - control ≥ 0). Reported p-values were adjusted using the single-step method. Model outcomes can be found in the supplementary materials (Table S2 and S3).

## Results

### Meta-analysis

Our meta-analysis provides a quantitative summary of the impact of microplastic soil contamination on major ecosystem process components, like soil aggregate formation, stabilization and organic matter loss (Fig. 2). We found strong limitations in the amount of available data for the three investigated effect sizes. The majority of studies reported effects of aggregate stability, while only one study presented data on soil organic matter decomposition. Microfibers made of polyester are the preferred microplastic type and polymer across studies, while only three studies tested polyacryl and a textile fiber mix, respectively. Particulate microplastics (e.g. fragments, beads, spheres) were targeted in five studies which tested a broad variety of polymers; with one study comparatively testing 6 different plastic polymers.

**Fig. 2.**
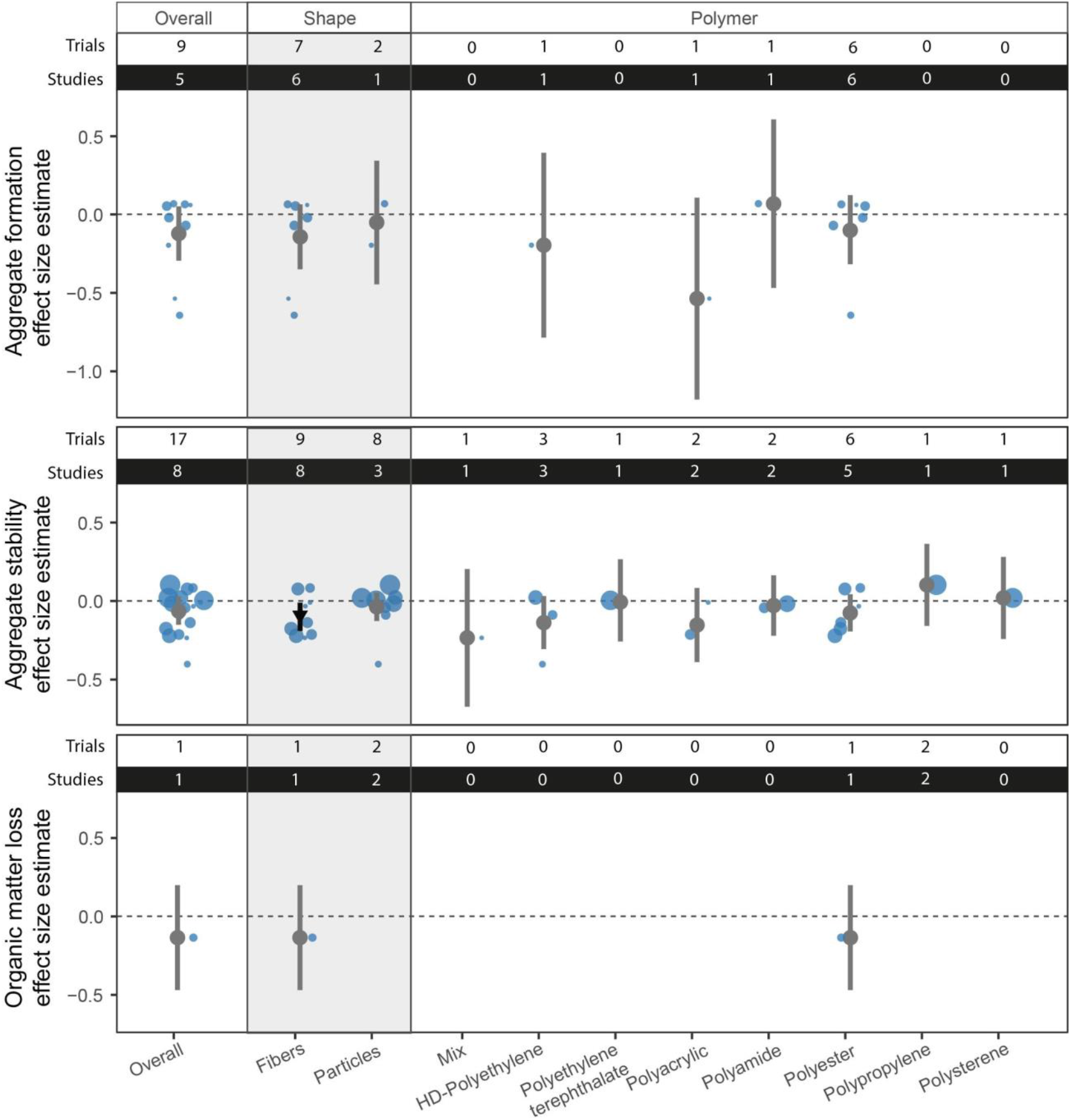
Overall, microplastic shape and polymer effects on soil aggregation process components and organic matter loss (decomposition of cellulose strips). Effect sizes and their variance are displayed as means and 95% confidence intervals. Effects are color-coded: neutral effect sizes are grey and negative effects are black with an arrow head pointing downwards; no positive effects were detected. Original data distribution is plotted in blue circles in the background, with circle size depicting study quality; studies with low variability and high number of replicates are represented with a large size. Number of experiments per study and the number of studies contributing to the presented effects are shown on top of each corresponding panel.

The magnitude of the three investigated effect sizes was overall neutral (the confidence interval includes zero). This could be ascribed to the limited available data and high variability among effect sizes values within groups, causing low statistical power. Only for polyester microfiber effects on aggregate stability, we detected a negative effect.

### Experiment

We tested 12 different plastic materials representing a combination of 4 different shapes (fibers, films, foams and fragments) and 8 distinct polymers (polyamide, polyester, polypropylene, polyethylene, polyethylene terephthalate, polyurethane, polycarbonate) for their impact on soil aggregate formation, stabilization and organic matter decomposition.

Across all our measured response variables the number of observed effects (negative or positive) for the different microplastic types were in the order films > fibers > fragments > foams.

The formation of new aggregates > 2 mm was affected most strongly among the measured parameters. We found that fibers consistently reduced this aggregate fraction irrespective of the plastic polymer (Fig. 3A, Fig. S3). For the other types, polyethylene and polyethylene terephthalate microplastics also reduced the amount of new formed aggregates; this finding was true across the different plastic types. The opposite was true for the polypropylene (PP) products (film – casted PP, foam – expandable PP, fragment – PP); for these materials we detected only neutral effects. Polyurethane foam and polycarbonate fragments yielded neutral and negative effects, respectively.

**Fig. 3.**
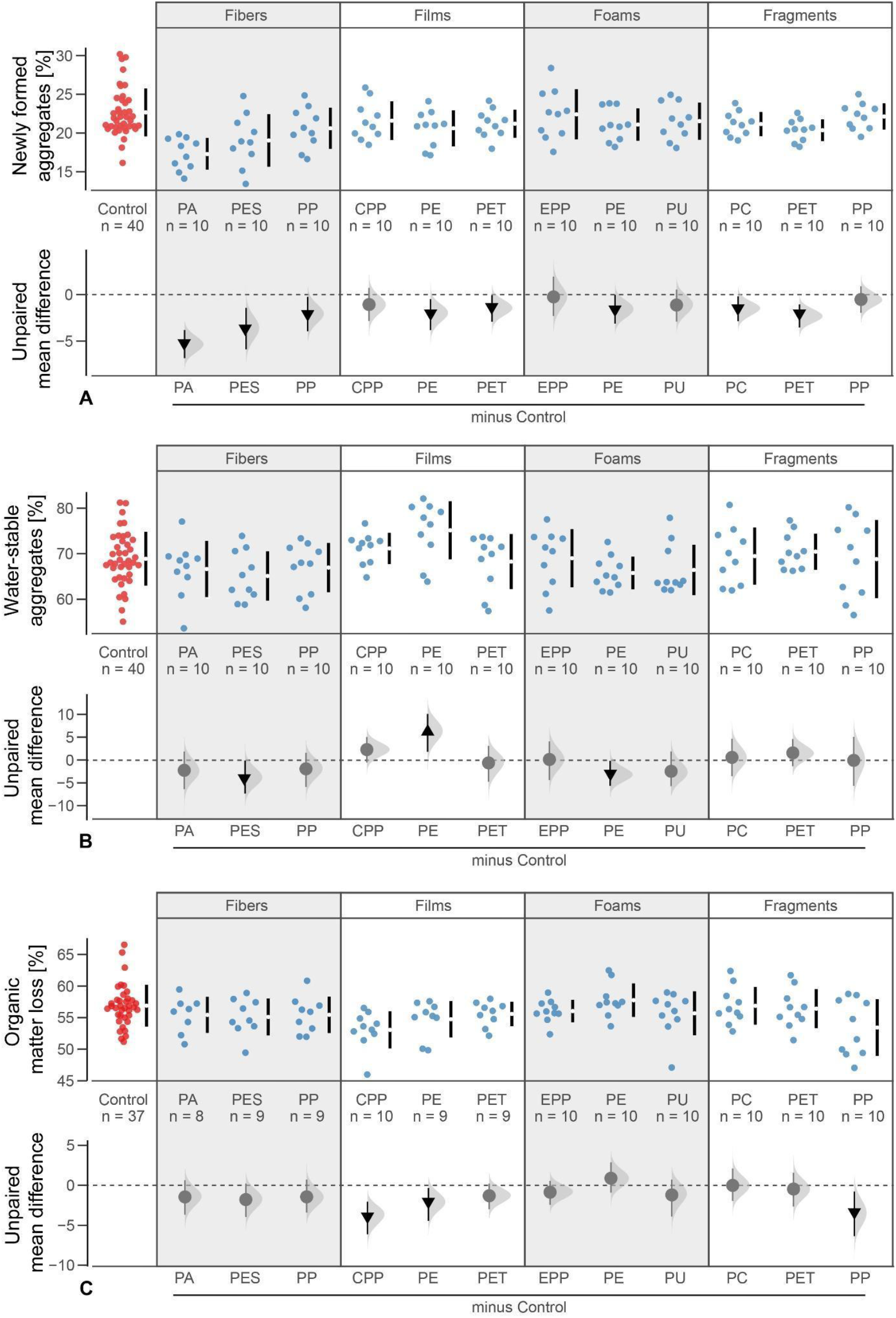
Microplastic shape and polymer effects on (A) newly formed aggregates > 2mm (in %), (B) stability of aggregates (in %) and (C) organic matter loss (in %). Raw data are presented as swarm plots in the first row of each panel and the number of replicates (n) are depicted. In the second row of each panel, multi-group estimation plots present the unpaired mean differences of the microplastic treatments and controls. The unpaired means (effect magnitude) are symbolized by circles and triangles and the corresponding confidence intervals (CIs; effect precision) by the vertical, black lines. The sampling error distribution is presented as a grey curve. Negative (arrow head pointing downwards) effects are colored in black whereas neutral effects (circle) are colored in grey. Outcomes of statistical analysis are presented in Table S2 and S3.

For aggregate stability, microplastic effects were less pronounced (Fig. 3B). Polyester fibers and polyethylene foams reduced, while polyethylene films increased aggregate stability. The other products had neutral effects.

Organic matter loss was the only variable not affected by microfibers (Fig. 3C). Instead, polypropylene films (CPP) and fragments and polyethylene films reduced the decomposition of the tea. All other products had neutral effects.

**Fig. 4.**
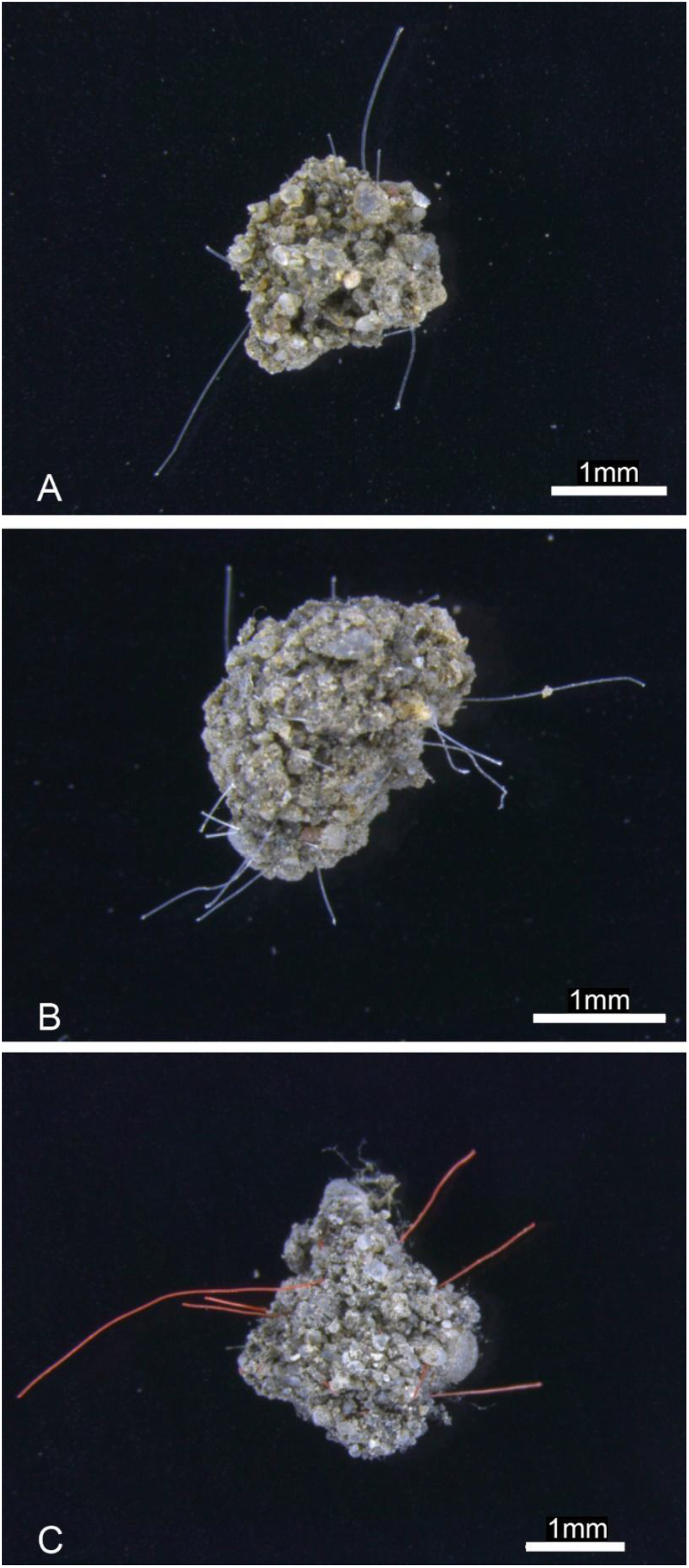
Photos of soil aggregates which formed during the course of the experiment. Aggregates incorporated microplastic fibers (polyamide fibers (A), polyester fibers (B), polypropylene fibers (C)).

## Discussion

### Meta-analysis

With this work we present insight into the importance of microplastic shape for soil responses, specifically soil structure and organic matter decomposition. Our meta-analysis revealed that polyester microfibers are the dominant plastic materials in experiments focusing on soil responses to microplastic contamination. For this material and shape, negative effects on soil aggregate stability are repeatedly reported (Machado et al., 2018; Boots et al., 2019; Liang et al., 2019; Machado et al., 2019; Zhang et al., 2019) which also manifested here in this analysis (Fig. 2). For particulated microplastics, a range of polymers has been tested, but with no clear pattern emerging for any of the investigated soil responses.

The data collection highlights the lack of a broad experimental base upon which to build mechanistic insights on the impact of microplastic – with its diverse shapes and chemistry - on major soil ecosystem process components. However, research on microplastic effects on terrestrial systems lags behind those of aquatic systems, hence we can expect a steady increase in new publications on the topic in the near future. Timely updates of these types of data syntheses are thus indispensable.

The focus on fibrous and particulate microplastics also reveals the complete absence of other shapes in the literature. There was no data available on effects of microplastic films or foams on the soil responses we investigated here. Experiments on the effects of film addition to a terrestrial system exist, but the product is always applied to the soil surface (e.g. Sintim et al., 2019), as is done during plastic mulching procedures, or in sizes exceeding the definition of microplastics. Thus it cannot be evaluated how microplastic films interact with the soil matrix, although they are widely distributed throughout different soils, including natural and agricultural systems (Brinton, 2005; He et al., 2018; Zhou et al., 2018; Ding et al., 2020; Qi et al., 2020).

A systematic investigation of different shapes is necessary to increase our understanding of the potential consequences of adding new materials with various shapes into soils. To fill this knowledge gap, we experimentally compared the effect of four different microplastic shapes (fibers, films, foams and fragments) on soil structure and organic matter loss.

### Experiment

#### Soil aggregation

Microfibers reduced the formation of new aggregates > 2 mm (Fig. 3A). There is no such clear pattern in the available literature, as demonstrated by our meta-analysis (Fig. 1).

However, there are experimental studies (Boots et al., 2019; Machado et al., 2019; Zhang et al., 2019) and field observations (Zhang and Liu, 2018) reporting that microfiber contamination led to reduced concentration of large aggregates. Our findings show clearly that this negative effect on aggregate formation is consistent across microfibers of different chemical composition (i.e. polymer), suggesting that this is indeed an effect primarily related to this linear shape. Observation of microplastic fibers sticking out of aggregates, into which they have apparently been integrated, supports the notion that microfibers introduce fracture points after being incorporated into new formed aggregates (see Fig. 4), likely facilitating breakdown when physical disturbance acts on the aggregate (Zhang and Liu, 2018; Rillig et al., 2019).

The detrimental effects of microfibers are potentially further modulated by the concentration (Machado et al., 2018) but also the average dimensions of the material. For both traits, we could not detect any robust pattern in our meta-analysis (data not shown). More trait information about applied microfiber material is needed to investigate the potential importance of e.g. curliness, flexibility or surface properties for aggregate formation.

In our experiment, microplastic films negatively affected aggregate formation while positively influencing aggregate stability (Fig. 1A and 1B). To our knowledge, these are the first data on microplastic film-induced effects on soil aggregation. Previous findings on plastic mulching residuals showed that microplastic films accelerate evaporation, decrease water content, but also reduce the soil tensile strength and bulk density while increasing soil porosity (Jiang et al., 2017; Wan et al., 2019). The microplastic films modify the soil matrix, introducing artificial pores and breaking planes, preventing the formation of large soil aggregates. The remaining smaller aggregates tended to be more stable. This might be due to the fact that our microplastic films had an average dimension of 4.4 mm; thus their size prevented their incorporation into smaller sized aggregates.

We found that microplastic foams tended to decrease the number of newly formed aggregates > 2 mm and the overall aggregate stability. Foams are clearly underrepresented shapes in microplastic research, although they are a ubiquitous and abundant contaminant, especially in coastal soils (Zhou et al., 2018), due to mariculture, tourism but also usage of insulating, construction, textile and packaging material (Shah et al., 2008). These light weight, porous, sponge-like structured shapes can soak with water or encrust with soil particles and soil microbes. As with films, the size will determine on which scale these materials will interact with the soil matrix. In our experiment, microplastic foams detrimentally affected the formation of aggregates. The negative impact on the stability on the remaining smaller aggregates could be attributed to potential additives (Gaylor et al., 2013).

The tested microplastic fragments showed a negative effect on the formation of new aggregates > 2 mm but had no detectable effect on aggregate stability. This negative impact is in line with the literature (Machado et al., 2018) but was not generally evident as shown by our meta-analysis (Fig. 1). Microplastic fragments had a range of soil responses from negative to positive. Fragments are less well incorporated into aggregates as compared to fibers (Zhang and Liu, 2018); in an agricultural soil, 72% were soil aggregate-associated while 28% dispersed in the soil. However, this process is probably determined by the fragment size. Other traits like surface smoothness/roughness, roundness/irregularity, rigidity/flexibility and brittleness/solidity potentially further modulate the induced soil responses.

Considering our results in the context of the shape dissimilarity hypothesis (Rillig et al., 2019), we here found strong evidence that microplastic fibers have a distinct negative impact on soil structure. Other shapes dissimilar to natural occurring shapes did not yield such obvious results. Fragments caused comparably detrimental effects as films, while for foams we detected most neutral responses. This may imply that beside shape, the material traits but even more the chemistry is important for the manifestation of microplastic-induced effects on soil structure.

This is supported by our finding that independently of the shape, polyethylene and polyethylene terephthalate reduced the amount of newly formed aggregates (Fig. 3A). This indicates that these polymers exhibited an inherent chemical property. Plastics can contain a variety of potential toxic substances (Hahladakis et al., 2018; Wang et al., 2019). Polyethylene can contain several additives like antioxidant agents (e.g. phenolics and phosphites), slip and blocking agents (fatty acids amides) or hydrophilic antistatic agents (e.g. polyethylene glycol esters) of up 0.01-0.5 % weight (Gachter and Muller, 1993). Migration of antioxidants and slip agents from low density polyethylene has been widely observed (within hours and days). Polyethylene terephthalate can contain plasticizers, UV-protectors, anti-static agents and post-consumer contaminants like limonene, benzaldehyde, anethole or benzophenone that have been absorbed by the plastic and can migrate from the plastic into the soil matrix (Widen et al., 2004; Hahladakis et al., 2018). The quantities of the migrating substances strongly depend on the hydrophobicity of the additive, the initial concentration and the characteristics of the surrounding environment (e.g. water, oil, solvents) (Li et al., 2016; Hahladakis et al., 2018). Due to their potential toxicity, the desorption of additives and their transition to the soil can affect soil processes such as microbial activity, subsequently affecting soil aggregation since microbial metabolites (e.g. exo-biopolymers like polysaccharides) can function as gluing substance and promote soil stability (Lehmann et al., 2017).

It is worth noting that microplastic effects resulting from laboratory studies might underestimate the true microplastic induced impact on the soil matrix compared to a field situation, because the microplastic is not constantly mixed with the soil. In the field, many processes cause constant mixing of microplastics in the soil matrix but also influence the soil aggregation process components formation and stabilization, such as plowing, bioturbation, dry-wet cycles and freeze-thaw cycles, the excretion of plant root exudates and entanglement by arbuscular mycorrhizal fungal hyphae. As a consequence, the microplastic is integrated into soil aggregates more thoroughly and thus the probability for potential impacts of microplastic increases (Zhang and Zhang, 2020).

#### Decomposition

In our experiment, microplastic films and fragments decreased the decomposition of organic matter (Fig. 3C); more specifically, the effects were detectable for casted polypropylene and polyethylene films and polypropylene fragments. The distinct microfiber impact found for soil aggregation did not manifest. In general, we found no evidence for a shape-but rather a polymer-induced effect.

The literature focusing on microplastic effects on soil organic carbon dynamics are scarce and when available only focusing on microplastic fibers and fragments, as demonstrated with our meta-analysis (Fig. 2C). The research findings so far suggest that microplasic effects on soil organic carbon dynamics are attributed to changes in soil physical parameters (i.e. porosity, connectivity, aeration) (Zhang and Zhang, 2020) and/or sorption and migration of chemicals and additives (Wang et al., 2016) which in turn affect soil microbial communities, their activity (Wang et al., 2016; Yang et al., 2018; Huang et al., 2019) and with them the mineralization rates of soil organic matter. For our study, only polyethylene microplastic films supported the notion of an important role of soil parameters, since it had negative effects on soil structure indicating a reduction in pore sizes and connectivity of microhabitats, which in turn could cause diminished microbial activity resulting in the detected reduction in organic matter loss. Microplastic films have a planar shape that could reduce the aeration of the soil, thereby reducing the activity of especially aerobic microbes.

For the polypropylene microplastic film and fragments, changes in soil structure could not be ascertained (Fig. 3). Hence it is questionable if additive migration from these materials might have induced the reductions in organic matter loss. However, additives are commonly detected in terrestrial samples (Whitacre, 2014) and it can be assumed that their migration potentially has toxic effects on microorganisms (Hahladakis et al., 2018) leading to impaired soil functions, such as decomposition.

The degradation of organic matter is an important ecosystem function and proxy for soil health that plays a crucial role in nutrient and especially carbon cycling (Lehmann and Kleber, 2015). If some microplastic types could disturb this process, it would have implications for soil C storage. To better understand this potential threat, we need more detailed studies on the influence of different microplastic types and polymers with exact information on their chemical properties.

## Conclusion and future research challenges

Our findings help to reduce the confusion about microplastic effects on soils, as it highlights the detrimental effect of microfibers on soil structure. Importantly, we revealed that other microplastic shapes affect soil structure as well. This effect depended on the polymer and should be the focus of further studies. However, there are still many unknowns in the equation, as information on polymer properties and additives is rare.

In summary, our meta-analysis and laboratory experiment in combination lend some support to the shape dissimilarity hypothesis, in particular in the case of fibers. However, other effects we observed appeared to be related to polymer type. More studies like this, dissecting particular aspects of microplastic diversity, are needed to achieve a more systematic and mechanistic understanding of the effects of these emerging pollutants.

## Acknowledgements

MCR acknowledges funding from an ERC Advanced Grant (694368) and the BMBF-funded project uPlastic. EFL acknowledges funding from the Deutsche Forschungsgemeinschaft (LE 859/1-1).

## Author contributions

A.L. designed the study. M.G. set up the experiment; M.G. provided experimental data; A.L. performed the statistical analysis; A.L., E.F.L., and M.C.R. wrote the manuscript; all authors contributed to the final version of the manuscript.

## Supplementary information

**Fig. S1.**
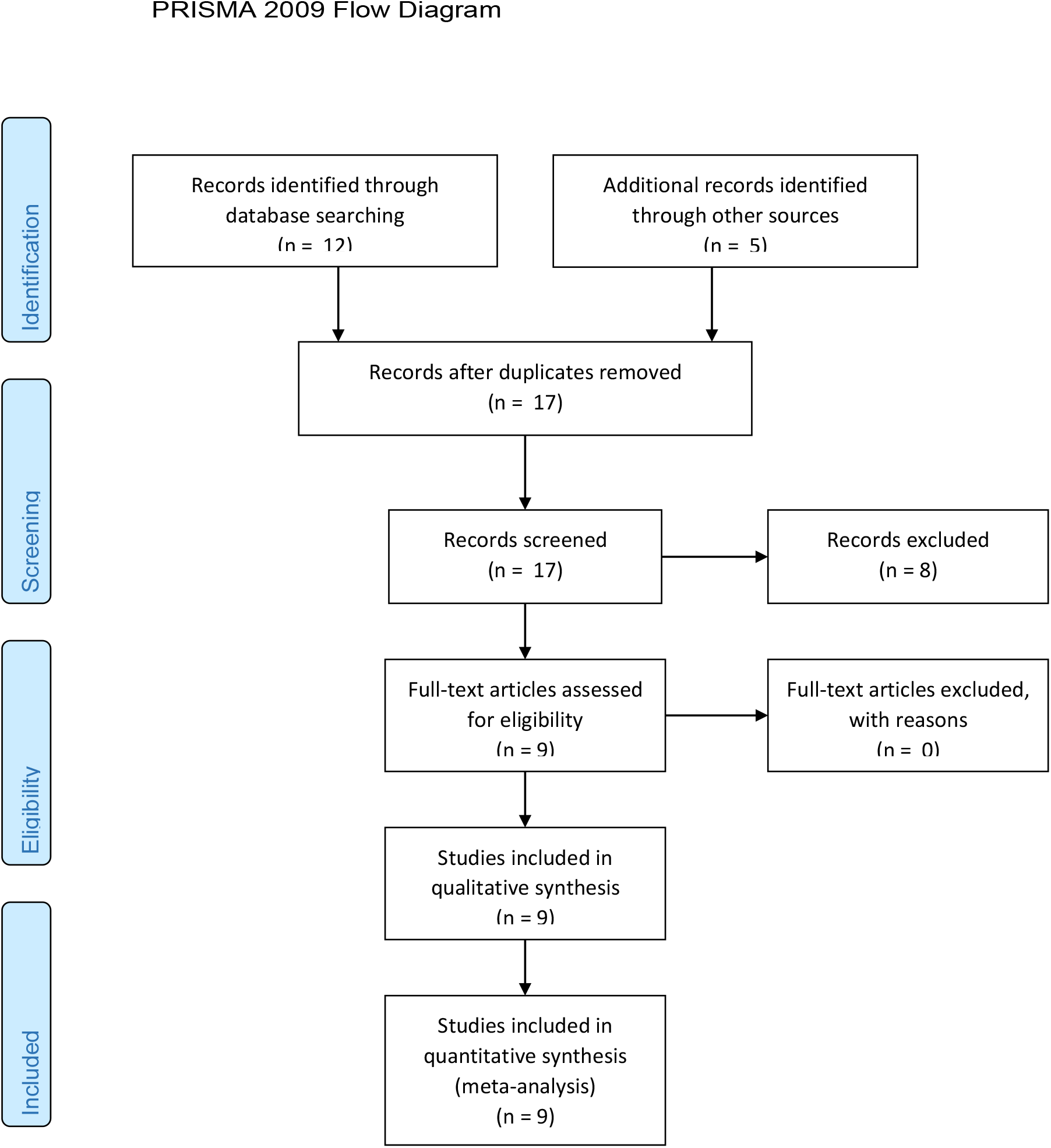
PRISMA flow diagram illustrating the process of data collection and quality control.

**Fig. S2.**
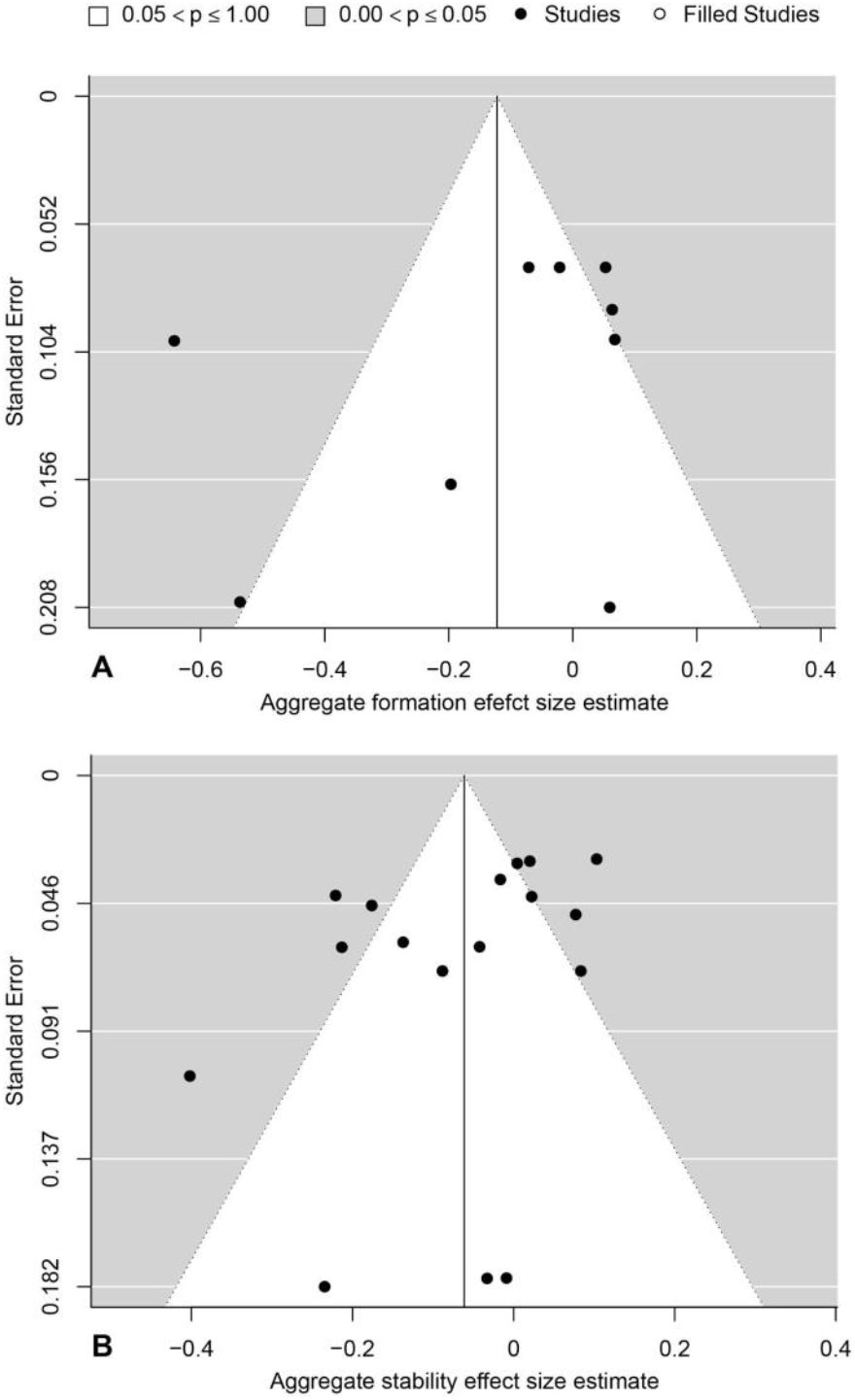
Funnel plots with trimfill option. Black dots represent effect size values from the corresponding aggregate formation, stabilization and soil Carbon datasets plotted against the model-derived standard errors. If a publication bias is detected, the trimfill option will include suggested (missing) effect size values (unfilled circles). This is not the case for our two testable datasets (namely aggregate formation and stability). For the soil organic matter loss dataset, no sensitivity could be applied since the dataset only comprised on effect size value. This suggests that we have no publication bias.

**Table S1.** Plastic products used to produce secondary microplastics.

| **Specific** |  |  |  |  |  |
| --- | --- | --- | --- | --- | --- |
| **Polymer** | **name** | **Full name** | **Type** | **Provider** | **Source** |
| PP |  | polypropylene | fiber | Hornbach (Art. 8442182, Seil 6cm x 2m) | https://www.hornbach.de/ |
| PA |  | polyamide synonymous to nylon | fiber | Hornbach (Art. 6702575, 0.35mm) | https://www.hornbach.de/ |
| PES |  | polyester | fiber | Hornbach (Art. Art. 8442173, Seil 8cm x 10m) | https://www.hornbach.de/ |
| PE | LDPE | polyethylene | foil | anti-algae folie (FolienBernhardt.de) | https://www.folien-bernhardt.de/produkte/folien |
| PET |  | polyethylene terephthalate | foil | oven bag, Toppits, EAN(s): 4008871209672 | from supermarket |
| PP | C/PP | cast polypropylene | foil | transparent folder; Stylex (Art. 40928) | from stationery shop |
| PET |  | polyethylene terephthalate | fragment | water bottle | from supermarket |
| PP |  | polypropylene | fragment | planting pots | from lab storage |
| PC |  | polycarbonate | fragment | CD, DVD | from supermarket |
| PU |  | polyurethan | foam | Hornbach (Art. 6411731) | https://www.hornbach.de/ |
| PE | LDPE | low density polyethylene | foam | package foam, HP laptop | from lab storage |
| PS | EPS | expanded polysterene | foam | transport box | from lab storage |

**Fig. S3.**
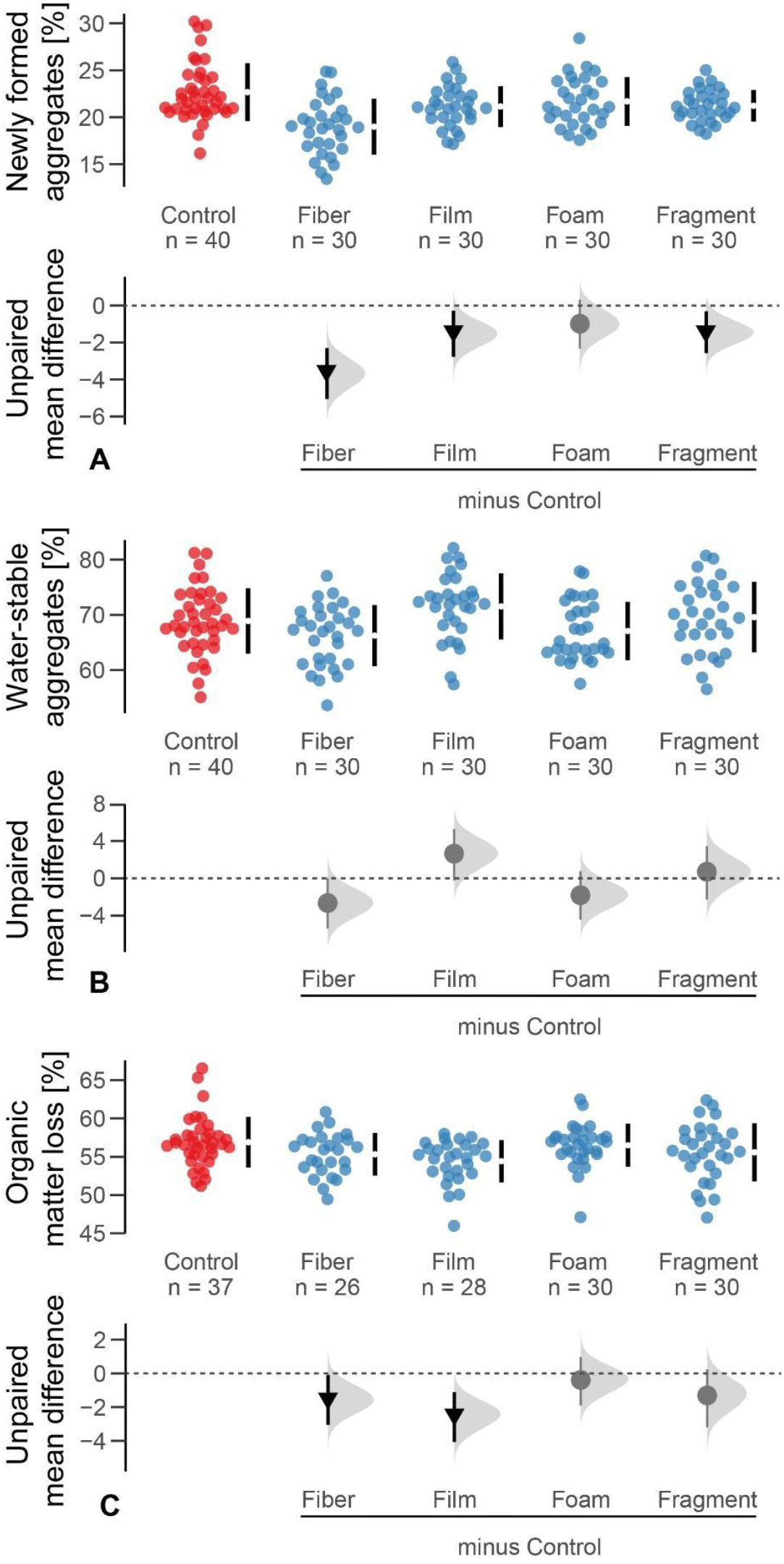
Microplastic shape effects on (A) newly formed aggregates > 2mm (in %), (B) stability of aggregates (in %) and (C) organic matter loss (in %). Raw data are presented as swarm plots in the first row of each panel and number of replicates (n) are depicted. In the second row of each panel, multi-group estimation plots present the unpaired mean differences of the microplastic treatments and controls. The unpaired means (effect magnitude) are symbolized by circles and triangles and the corresponding confidence intervals (CIs; effect precision) by the vertical, black lines. The sampling error distribution is presented as a grey curve. Negative (arrow head down) effects are colored in black whereas neutral effects (circle) are colored in grey. Outcomes of statistical analysis are presented in Table S2 and S3.

**Table S2.**
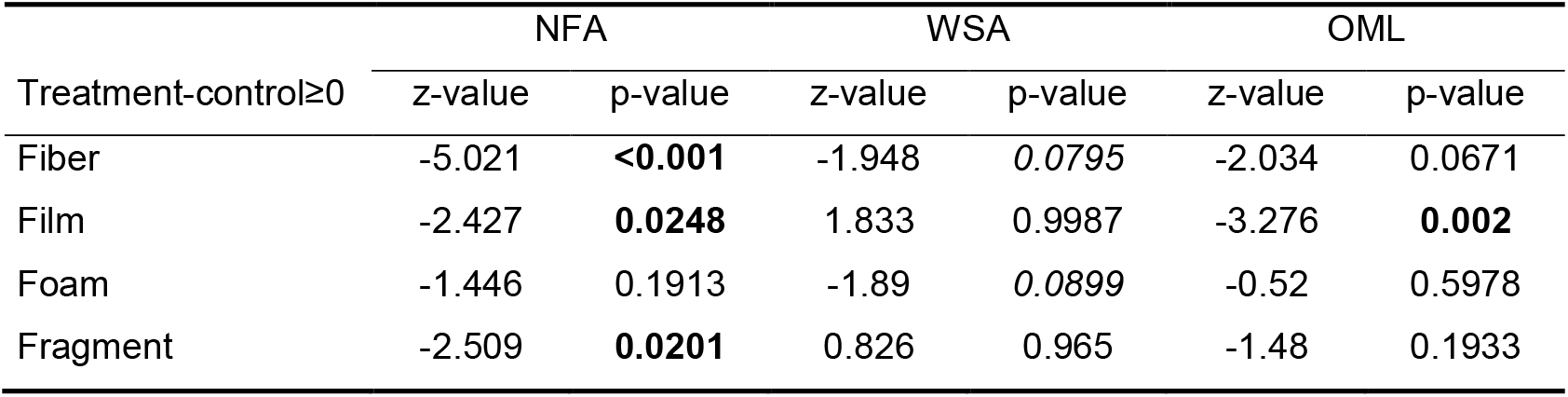
Outcomes for newly formed aggregates (NFA in %, n=70), water-stability of aggregates (WSA in %, n=70) and organic matter loss (OML in %, n=61) and microplastic shapes. A p-value < 0.05 was considered significant and marked in bold.

**Table S3.** Outcomes for newly formed aggregates (NFA in %, n=70), water-stability of aggregates (WSA in %, n=70) and organic matter loss (OML in %, n=61) and microplastic polymer per shape. A p-value < 0.05 was considered significant and marked in bold.

| Treatment - control >= 0 |  | NFA |  | WSA |  | OML |  |
| --- | --- | --- | --- | --- | --- | --- | --- |
|  |  | z-value | p-value | z-value | p-value | z-value | p-value |
| Fiber | PA | -6.577 | **<0.0001** | -1.04 | 0.709 | -1.27 | 0.56571 |
|  | PES | -3.081 | **0.01123** | -1.939 | 0.225 | -1.605 | 0.37739 |
|  | PP | -2.118 | 0.14265 | -0.989 | 0.735 | -1.306 | 0.5448 |
| Film | CPP | -1.161 | 0.58602 | 1.603 | 1 | -1.939 | 0.21859 |
|  | PES | -2.339 | 0.0878 | 2.824 | 1 | -3.548 | **0.00226** |
|  | PET | -1.939 | 0.20226 | -0.29 | 0.957 | -1.541 | 0.41187 |
| Foam | EPP | -0.22 | 0.93908 | 0.061 | 0.989 | -1.09 | 0.66493 |
|  | PE | -1.91 | 0.21316 | -2.119 | 0.158 | 0.886 | 0.99974 |
|  | PU | -1.251 | 0.53727 | -1.237 | 0.602 | -0.975 | 0.72407 |
| Fragment | PC | -2.176 | 0.12631 | 0.272 | 0.996 | -0.004 | 0.98054 |
|  | PET | -3.481 | **0.00287** | 1.012 | 1 | -0.41 | 0.92538 |
|  | PP | -0.728 | 0.79465 | -0.019 | 0.984 | -2.272 | 0.10966 |

